# Retronasal Habituation: Characterization and Impact on Flavor Perception using Time-Intensity

**DOI:** 10.1101/274514

**Authors:** Robert Pellegrino, Addison Atchley, Simrah Ali, Joel Shingleton, Curtis R. Luckett

**Author notes:** Corresponding author: Curtis R. Luckett; Department of Food Science, University of Tennessee, 2510 River Drive, Knoxville, TN 37996, U.S.

## Abstract

Olfactory habituation results from prolonged exposure to an odor, leading to perceptual changes defined by several characteristics. To date, human habituation research has focused on orthonasal olfaction which is perceived externally while ignoring internal routes of odor perception related to flavor. In our study, we conducted two experiments to characterize retronasal olfactory habituation and measured its impact on flavor perception. In Experiment 1, 22 participants rhythmically breathed a food odor (lime), non-food odor (lavender), and blank (propylene glycol) that was presented using an orally-adhered strip, while rating the odor intensity using the time-intensity procedure. After a 10-minute exposure, the participants ate a lime-flavored gummy and rated the lime flavor. In Experiment 2, the same procedure was performed for a low-level lime odor, a simple (lime oil) and complex (lime oil + sucrose + citric acid) beverage as the flavor stimuli. Our results demonstrated two known principles of habituation for retronasally presented odors: 1) prolonged exposure lead to decreased perception, 2) weaker stimuli lead to more rapid habituation. Additionally, we found that the non-food odor habituated slower than the food odor; however, the participants seemed to recover simultaneously upon food and beverage consumption leading to no change in flavor perception.

## Introduction

Humans perceive odors via two presentation routes, orthonasal and retronasal. The orthonasal route is characterized by external detection of odor through the nostrils. Retronasal refers to odors detected internally, in which odors are brought into the nose from the mouth through the nasopharynx (Mozell et al., 1969). Although the same odor may present itself through both routes, each mode of presentation interprets the odor differently at the receptor and central levels (Bojanowski and Hummel, 2012). For instance, flow paths change among anatomical differences and sorption rates of odorants vary across the olfactory mucosa leading to a range of activation patterns along the olfactory epithelium between routes (Yang et al., 2007; Scott et al., 2014).

A well-known phenomenon of flavor perception is the referral of odors to the mouth when in reality pulses of odors during the breakdown and swallowing of food are detected retronasally. Food odors are typically associated with concurrent taste stimulation, creating associative learning that leads to the distinctive central processing of retronasally presented food odors. Retronasal food odors activate areas of the brain related to the mouth and overlap with taste-specific regions such as the insula (Small et al., 2004, 2005). Considering taste and smell interactions in flavor perception, retronasally-presented food odors are often associated with concurrent taste stimulation leading them to take on “taste-like” properties (Lim and Johnson 2012; Stevenson, Prescott, and Boakes 1999; Stevenson, Boakes, and Prescott 1998). When odors are presented retronasally they are perceived as less intense than an orthonasal mode of presentation (Hummel et al., 2016). However, little is known about basic habituation properties of retronasally perceived odors (Pellegrino et al., 2017). Attempts to characterize retronasal olfactory habituation have been relatively unsuccessful (Pierce and Simons, 2018). It has been suggested that additional studies, using more naturalistic delivery methods are needed to better understand retronasal olfactory habituation (Pierce and Simons, 2018).

Habituation refers to the decrease in *behavioral response* as a result of repeated and prolonged stimulation, while adaptation refers to the *physiological processes* (peripheral and central) that constitute a change in this response. Historically, there has been some confusion over these terms, which has led to them being used interchangeably. Clouding the difference even more, the two phenomena are extremely difficult to differentiate through psychophysical measurements. In their seminal paper on habituation, Thompson and Spender (1966) outlined 9 key principles of habituation, and these were revisited by Rankin et al. (2009). Specific to human olfactory, these principles of habituation are measured with a subjective reduction of intensity (perception of the strength of the odor) while direct reductions in underlying neuronal processes (pre-and post-glomerular) are termed adaptation and measured with physiological techniques [e.g. electro-olfactogram (EOG), electroencephalography (EEG), etc.]. All senses undergo adaptation that modifies the perception of the stimuli that may lead to behavioral changes defined by habituation (Thompson and Spencer, 1966; Rankin et al., 2010).

Odors are usually presented in plumes representing several different stimuli (or parts of a stimuli) and habituation helps segment them. This rapid segmentation allows us to separate out informative odors from non-informative ones and novel odors from those that are already present (Kadohisa and Wilson, 2006; Uchida et al., 2006; Linster et al., 2007; Gottfried, 2010). This type of orthonasal habituation has been well studied and characterized in humans, but retronasal odors (often perceived while eating) have received less attention (Dalton, 2000; Pellegrino et al., 2017). Similarly, little is known about the impact retronasal habituation has on flavor in a natural breathing paradigm.

Two experiments were conducted to characterize principles of retronasal olfactory habituation and measure its impact on flavor perception. Specifically, two of the nine principles that have been demonstrated in orthonasal olfactory habituation were characterized: Principle 1) repeated applications of stimulus result in decreased responses and Principle 5) weaker stimuli lead to more rapid habituation (Thompson and Spencer, 1966; Rankin et al., 2010). The first experiment was designed to determine habituation rates of a food (lime) and non-food odor (lavender) compared to a control (propylene glycol). Habituation was induced with an odor-containing strip adhered to the roof of the mouth and rhythmic breathing (inhaling through the mouth and exhaling out the nose). Following habituation, the impact of a flavor was measured on a lime gummy. In a follow-up experiment, the authors repeated the same procedure from Study 1 with a lower intensity of lime odorant. After habituation, participants were asked to rate the flavor intensity of two lime beverages, one simple (lime oil + water) and one complex (water + lime oil + sucrose/citric acid).

## Experiment 1

### Materials and Methods

#### Participants

Twenty-six volunteers were recruited for this study. Participants reported a good sense of smell, had no allergies or food restrictions and were not pregnant or smokers. Each participant was measured on their ability to identify odors using Sniffin’ Sticks, and only those who correctly identified 75% or higher of the odors were asked to complete the study (Hummel et al., 1997). Olfactory ability screening eliminated 3 individuals and 1 subject dropped out of the study, leaving 22 (12 women) with an age ranging from 22 to 64 (M ± SD, 33.55 ± 11.08).

Participants were asked to refrain from eating or drinking at least one hour before the experiment, and asked to exclude menthol products from their diet on the day of testing. All participants signed an informed consent and were compensated for their time spent participating. This experiment (and the follow-up protocol) were conducted according to the Declaration of Helsinki for studies on human subjects and approved by the University of Tennessee IRB review for research involving human subjects (IRB #16-03417-XP).

#### Stimuli and retronasal strips

Two odorants were used as the main stimuli, lime (0110-0500, LorAnn Oils and Flavors, Lansing, MI) and lavender (2270-0500, LorAnn Oils and Flavors, Lansing, MI). Propylene glycol was used to dilute the pure odor stimuli and as a control. The lime solution (diluted with propylene glycol) was made at a concentration of 33.3% (v/v) while the lavender solution was made at a concentration of 26.6% (v/v). These concentrations were based on a pilot study, in which individuals (N=10) rated a drop (10 μL) of several odor concentrations on 2 X 2 cm filter paper with a 150-point line scale for pleasantness (“Extremely Unpleasant” to “Extremely Pleasant”) and intensity (“Not intense at all” to “Extremely Intense”). The chosen concentrations were not significantly different in either property. The lavender and lime solutions were rated 10.8 and 9.3 respectively. For pleasantness lavender and lime solutions were rated 9.7 and 9.4, respectively.

In order to adhere the odorant delivery system to the roof of the mouth, Pullulan (Hayashibara, Okayama, Japan) was utilized. Pullulan is a linear homopolysaccaride of glucose and is synthesized from the fungus *Aureobasidum pullulans.* It has a variety of distinctive traits that include unique linkage patterns allowing for creation of strong impermeable films that are translucent, tasteless and edible (Leathers, 2003). Additionally, upon hydration pullulan films form an adhesive gel. The adhesive gel, formed upon contacting saliva, is vital to the adherence of the odor stimuli in this paper. To make the strips, an aqueous solution (100 mL) of pullulan (5% w/v) dissolved in distilled water was cast on a sanitized tray (215.9 X 279.9 X 12.7 mm) and allowed to evaporate at room temperature (23 °C) for 24 hours. After the film formation, a spritz bottle with distilled water was used to moisturize the surface and a paper composed of cotton linters was added to the surface. This film was cut into 10 X 20 mm rectangular strips and stored into a snap-sealed bag at room temperature. Five minutes before use in the experiment, 10μL of odor solution (either lime or lavender) or propylene glycol was added to the middle of the strip on the side composed of cotton linters. Figure 1 depicts the odor stimulus and its application to the hard palate. The strip was then immediately placed into a small, plastic container (30 mL) with a lid.

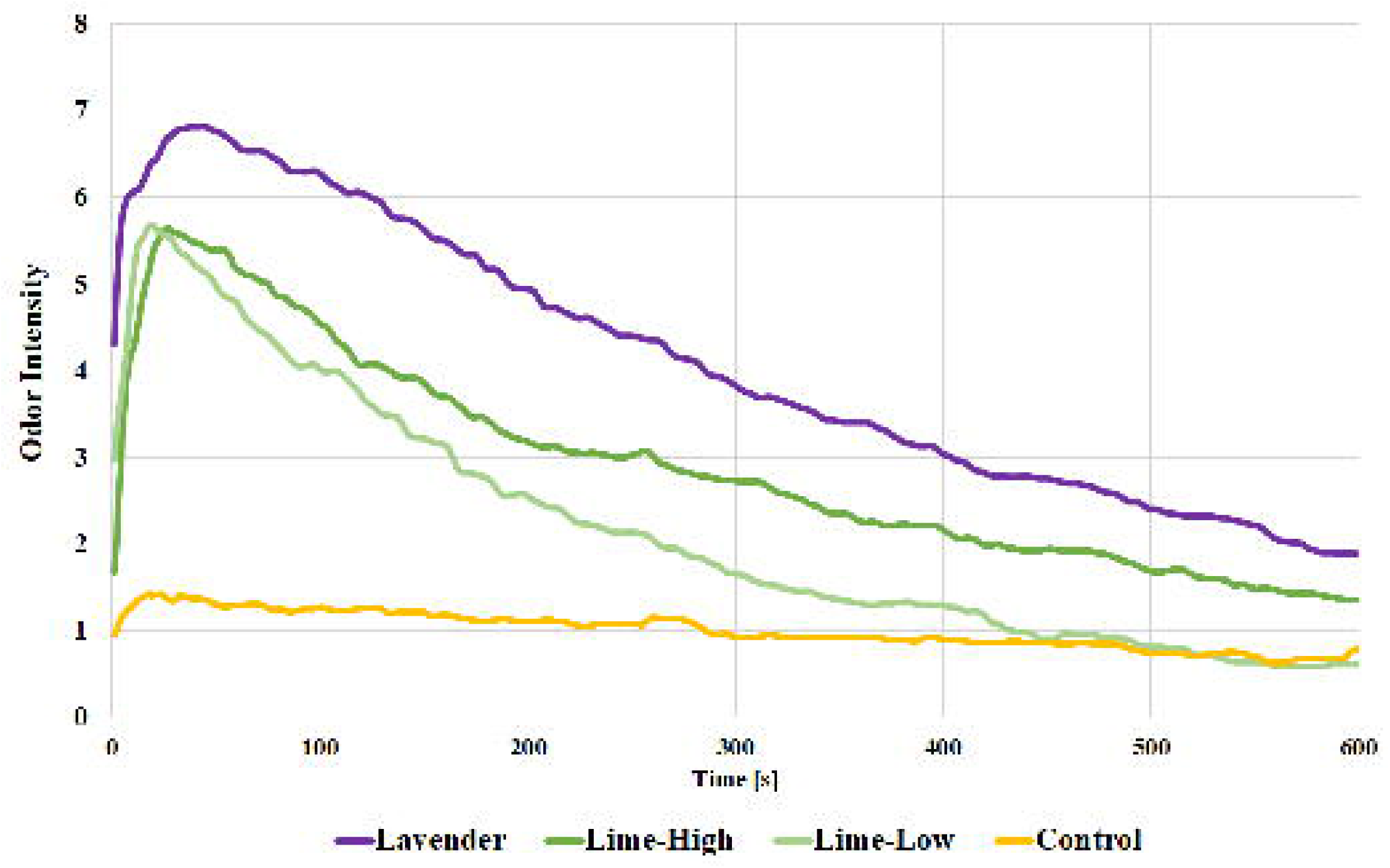

To create the flavor stimuli, sucrose (Domino Foods Younkers, NY), glucose syrup (Caulet, Erquinghem-Lys, France), sorbitol (4mular, Irvine, CA), and citric acid (SAFC, Switzerland) were mixed together and heated using a double boiling system until forming a homogenous solution (see Table 1). Gelatin (250 bloom, Knox, E.D. Smith Foods, Winona, Canada) was dissolved separately in boiling water, then added to the solution. The solution was brought to room temperature (23 °C) and lime oil (6 μL of per gummy; LorAnn Oils and Flavors, Lansing, MI) was incorporated. The solution was then poured into corn starch-dusted, hemi-spherical silicone molds (11.2 cm^3^) and allowed to harden in a refrigerator (4°C) overnight.

**Table 1.** List of ingredients and their amount used to make lime gummies.

| **Ingredient** | **Amount (g)** |
| --- | --- |
| Gelatin | 1.20 |
| Water | 3.10 |
| Glucose Syrup | 6.00 |
| Sucrose | 3.00 |
| Sorbitol | 0.30 |
| Citric Acid | 0.03 |

#### Time-Intensity (TI) task

To measure the temporal dynamics of retronasal odor habituation, time–intensity (TI) analysis (Larson-Powers and Pangborn, 1978) was used with the sensory analysis software, Red Jade (Redwood City, CA). Overall odor intensity of the strip was rated on a 10-point horizontal scale, representing overall odor intensity. The scale was anchored by “Maximum” on the right and “0” on the left via the TI scaling software; the sampling rate was 0.5 s. The TI parameters used, as well as their definitions, are presented in Table 2.

**Table 2.**
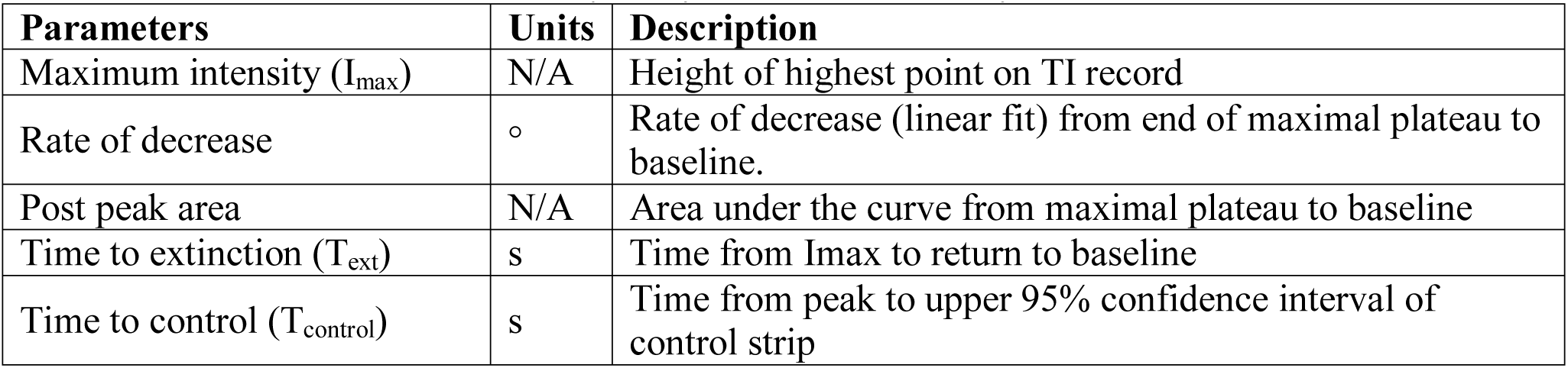
Parameters of Time-Intensity analysis used in this study

| **Parameters** | **Units** | **Description** |
| --- | --- | --- |
| Maximum intensity ($I_{\max}$) | N/A | Height of highest point on TI record |
| Rate of decrease | ° | Rate of decrease (linear fit) from end of maximal plateau to baseline. |
| Post peak area | N/A | Area under the curve from maximal plateau to baseline |
| Time to extinction ($T_{\text{ext}}$) | s | Time from $I_{\max}$ to return to baseline |
| Time to control ($T_{\text{control}}$) | s | Time from peak to upper 95% confidence interval of control strip |

#### Procedure

Prior to beginning the experiment, individuals were briefed on time-intensity, scale usage and the test procedure. Additionally, they were familiarized with a blank strip, how to insert it into the mouth (pullulan side adheres to the roof of the mouth), trained on their breathing pattern (inhale through the mouth, exhale through the nose) with a metronome (at 40 bpm), and asked not to touch the sample with their tongue. Next, they were positioned in an off-white, noise-controlled booth, and given an oyster cracker to practice the TI task. For the cracker, the TI task was to rate the intensity of salt over the 30 s of mastication. Following the practice and a water rinse, they were given two closed plastic containers (30 mL), one with a retronasal odor strip (odorized or blank) and one with a lime gummy for the main TI task. They were then instructed to open the strip container, adhere the strip to the roof of their mouth, and start the TI task once the odor strip was in place. The TI task was set for 10 min and individuals were encouraged to breath, as trained, the entirety of the procedure even if there was no perception of an odor.

Immediately after the task, individuals were instructed to remove the strip, eat the gummy and rate the intensity of the lime flavor on a 150-point line scale from “Not intense at all” to “Extremely intense”. Participants were given a 5-min break to fully recover from habituation and then this procedure was repeated for another strip condition. Participants were tested twice for each of the three strip conditions (blank, lavender and lime) which were randomly distributed over three days.

#### Statistical Analysis

Time-intensity curves for each condition were created from the intensity rating for each second of the 600 second long trail (10 min) and parameters were extracted from these curves based on equations set by ASTM international (ASTM International, 2017). Parameters of interest were I_max_ (or maximum intensity), post peak area, rate of decrease, and T_ext_ (or extinction time) (see Table 2 for description of each). However, many individuals never reached a point of complete cessation of smell [extinction value of zero, see (de Wijk, 1989)] and thus had a 600 s duration time or no extinction time. This was true for each odor and even the blank condition. Therefore, a modified T_ext_ parameter was created called T_control_. This value was the time at which the intensity of the stimuli was less than the upper 95% confidence interval of the blank odor strip intensity. All parameters were compared across odor strip conditions with a repeated measures ANOVA except T_control_ which used a paired T-test. Pairwise post-hoc comparisons were performed using Tukey’s HSD test.

Individuals who did not show reliable flavor intensity scores were excluded from the analysis. Four individuals (3 females) were removed from analysis due to unreliable results (variation of over 1/3 of the scale between replications) leaving a total of 18 (9 woman) ranging in age from 22 to 64 (32.67 ± 11.12) to be analyzed. The flavor intensity scores were set as the response of a repeated measures ANOVA looking at differences among odor strip conditions. All analysis was done in JMP (version 13.0; SAS Institute, Cary, NC, USA), with a statistically significant difference defined as p < 0.05.

## Results

### Time-Intensity Parameters

The I_max_ (or maximum intensity) among the strips was significantly different [F(2, 34) = 47.1898, p < 0.001] with both odor strips [lime (M ± SD = 6.36 ± 1.88) and lavender (7.42 ± 1.76)] having a larger I_max_ than the control (M ± SD = 3.00 ± 2.37, p < 0.001), but there was no difference between the two odors (p = 0.082). The post peak area was also significantly different among strip conditions [F(2, 34) = 37.56 p < 0.001). Post-hoc analysis (Figure 2) shows that the control had a smaller post peak area than both odor conditions (p < 0.001) and lavender had a larger post peak area than lime (p = 0.01). Lastly, T_ext_ was significantly shorter for the control than both odors [F(2, 34) =34.7616, p < 0.001], but did not differ among the lavender and lime. However, T_control_ revealed the lime odor to have a shorter extinction time than the lavender odor condition [T(1, 17) = -2.41, p = 0.03]. There was no difference in the rate of decrease between the strip conditions [F(2, 34) = 2.1320, p = 0.13]. All means and significant differences are shown in Table 3.

**Table 3.** Time-Intensity (TI) parameters for oral strip conditions.

| **Strip Condition** | **Time-Intensity Parameters** |  |  |  |  |
| --- | --- | --- | --- | --- | --- |
| | $I_{\max}$ | Post peak area | $T_{\text{ext}}$ | Rate of decrease | $T_{\text{control}}$ |
| Control | $3.00 \pm 2.37$ b | $468.42 \pm 604.78$ c | $266.17 \pm 257.58$ b | $-0.04 \pm 0.09$ a | - |
| Lavender | $7.42 \pm 1.76$ a | $2292.94 \pm 1022.87$ a | $572.94 \pm 50.45$ a | $-0.01 \pm 0.01$ a | $461.47 \pm 143.90$ a |
| Lime High | $6.36 \pm 1.88$ a | $1640.31 \pm 925.45$ b | $500.25 \pm 125.85$ a | $-0.01 \pm 0.01$ a | $386.28 \pm 177.73$ b |
| Lime Low | $6.18 \pm 1.77$ | $1253.89 \pm 549.08$ * | $455.5 \pm 162.06$ | $-0.02 \pm 0.02$ * | $298.74 \pm 140.3$ * |

Note. Means with differing subscripts within TI columns are significantly different at the adjusted $p < 0.05$ based off Tukey HSD. \* signifies $p < 0.05$ between lime low and high.

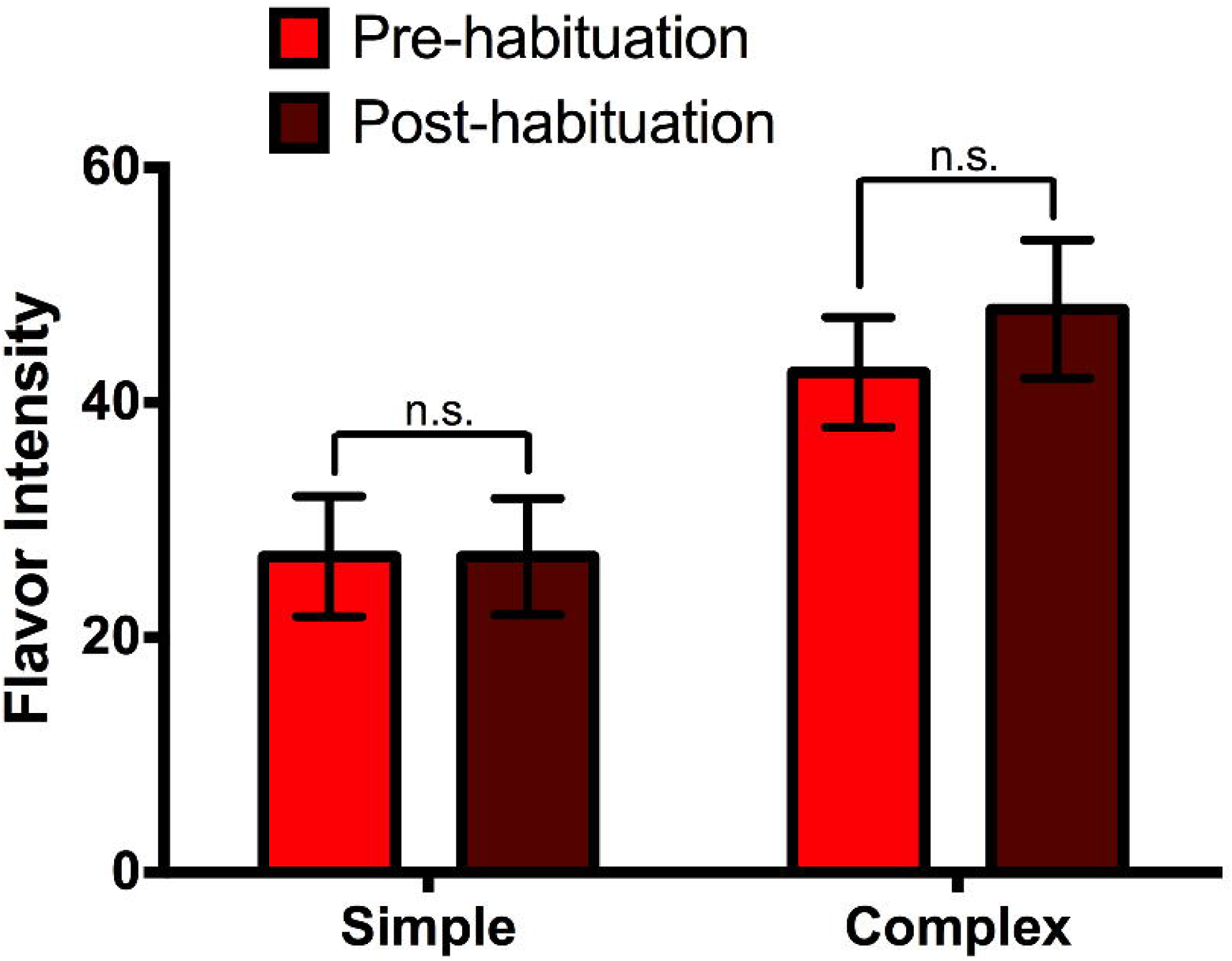

### Habituation Impact of Flavor

No overall difference of flavor intensity existed between the strip condition [F(2, 34) = 1.43, p = 0.25]: lime (7.96 ± 4.24), control (8.76 ± 3.01) and lavender (8.91 ± 3.66).

## Experiment 2

To examine another primary principle to habituation (principle 5), this study was designed to estimate the habituation curve of the lime odor at a lower concentration. Additionally, to further understand the impact of habituation on flavor (and cover limitations from Experiment 1), a simple and complex beverage was served and compared against a control using a logarithmic intensity scale.

### Participants

Nineteen individuals (9 women) with an age ranging from 20 to 64 years old (33.58 ± 11.88 years) participated in this study. Participants reported a good sense of smell, had no allergies or food restrictions and were not pregnant or smokers. Similar to Experiment 1, only those who correctly identified at least 75% of the odors were invited to participate in the study. Participants were asked to refrain from eating or drinking at least one hour before the experiment, and additionally asked to exclude menthol products from their diet on the day of testing. All participants signed an informed consent and were compensated for their time.

### Stimuli

A 10 % (v/v) solution of lime oil (0110-0500, LorAnn Oils and Flavors, Lansing, MI) dissolved in propylene glycol was prepared for this experiment. This concentration of lime odor was determined with a preliminary study (N = 10) showing a significantly lower intensity score compared to the odor concentration from Experiment 1, when perceived orthonasally, and rated on a 150-point scale (“Not intense at all” to “Extremely Intense”). The lower concentration lime odor was rated a 5.3 and a 10.2 for intensity and pleasantness, respectively. Exactly as in Experiment 1, 10 μL of the lime solution was added to a pullulan-based strip.

Instead of gummies, a beverage was used in this experiment. A beverage was chosen because there is less mechanical stimulation during drinking than eating. Two stimuli were made to represent a simple (without taste) and complex (with taste) beverage. The complex beverage contained sucrose (7%, w/v), citric acid (0.08%, w/v) and lime oil (0.00006%, v/v) dissolved in distilled water while the simple beverage contained only lime oil at the same concentration dissolved in water. Beverages were presented in a sealed plastic container (60 mL) with 50 mm long straws protruding from a hole placed in the top (another hole was placed at the far end to relieve pressure). This presentation reduced possible orthonasal dishabituation/compensation when bringing the drink up to the face for consumption.

### Procedure

A similar procedure to Experiment 1 was used in this follow-up experiment (including the practice TI task). However, before participants placed the odor strip in their mouth they were given either a simple or complex beverage. They were asked to drink the beverage and rate the intensity of lime flavor on a labeled estimation scale – a semantic scale measuring intensity with quasi-logarithms spacing of verbal labels (Green et al., 1996). Next, they were given two containers, one with an odor strip (at low concentration only) and the same beverage type. After completing the TI task (as described in Experiment 1), the participants were asked to immediately drink the beverage and rate the intensity of lime flavor with the same type of scale. The participants were then given a 5 min break to help recover their sense of smell, and this procedure was repeated for the other type of beverage (either simple or complex). The order of beverage type was randomly distributed among participants.

### Statistical Analysis

The time-intensity curve for each beverage type was calculated from the intensity rating per second of the 600 second trail (10 min). As the habituation procedure was identical to that of Experiment 1, the TI parameters were extracted from a pooled curve. The T_control_ was calculated for the low-lime condition. This parameter and the other parameters from Experiment 1 were compared for low-lime and high-lime odor strip conditions using an independent t-test.

The effect of habituation on flavor intensity was compared with two dependent t-tests measuring flavor intensity differences between consumption of the stimuli pre-and post-habituation for complex and simple conditions. Additionally, a dependent t-test compared simple and complex flavor intensity differences in pre-post consumption to see if one had more impact than the other.

All analysis was done in JMP (version 13.0; SAS Institute, 2000, Cary, NC, USA) with a statistically significant difference defined as p < 0.05.

## Results

### Time-Intensity Parameters

As shown in Table 3, high lime had a significantly larger post peak area than low lime strip condition, T(1, 72) = 2.20, p = 0.03. Similarly, there was a faster rate of decreasing slope for low lime compared to the high lime, T(1, 72) = 2.38, p = 0.02. There was no difference in I_max_ [T(1, 72) = 0.41, p = 0.68] and T_ext_ [T(1, 72) = 1.32, p = 0.19) between the two conditions; however, T_control_ revealed a shorter extinction time for low lime compared to high lime [T(1, 72) = 2.36, p = 0.02]. All TI curves (from Experiment 1 and 2 strip conditions) are overlaid in Figure 3 to visually depict differences.

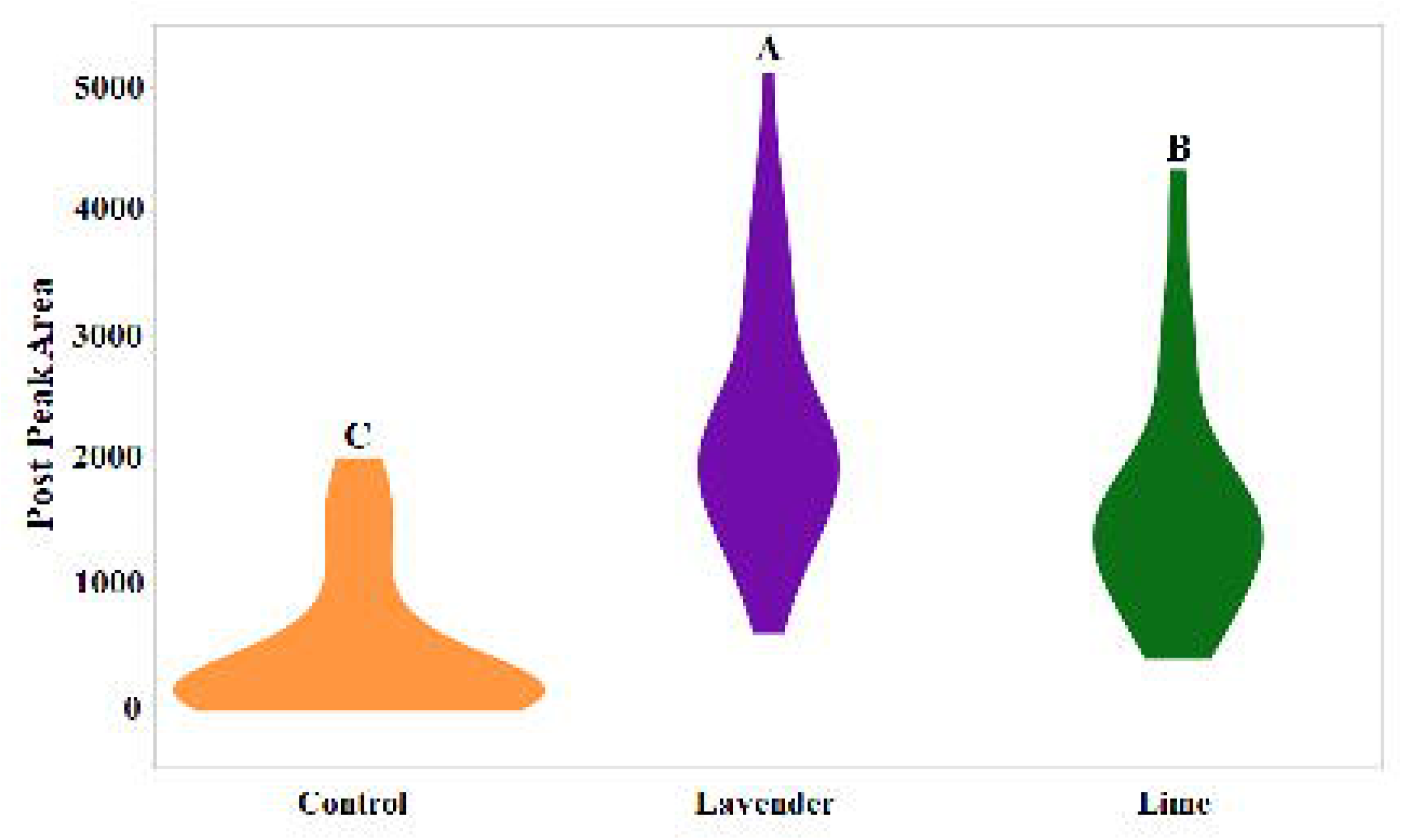

### Habituation Impact on Flavor

As observed in Figure 4, there was little effect of habituation on beverage flavor. There was no difference in flavor intensity before and after habituation of the low lime odor strip for the simple [T(1, 18) = 0, p = 1) and complex beverages [T(1, 19) = 1.27, p = 0.22]. Furthermore, the differences between pre-post did not vary among complex and simple beverages [T(1,18) = 1.02, p = 0.32]; however, the control complex beverage (42.57 ± 20.36) was rated as having a more intense lime flavor than the control simple beverage (26.89 ± 22.32) [T(1, 18) = 2.24, p = 0.04].

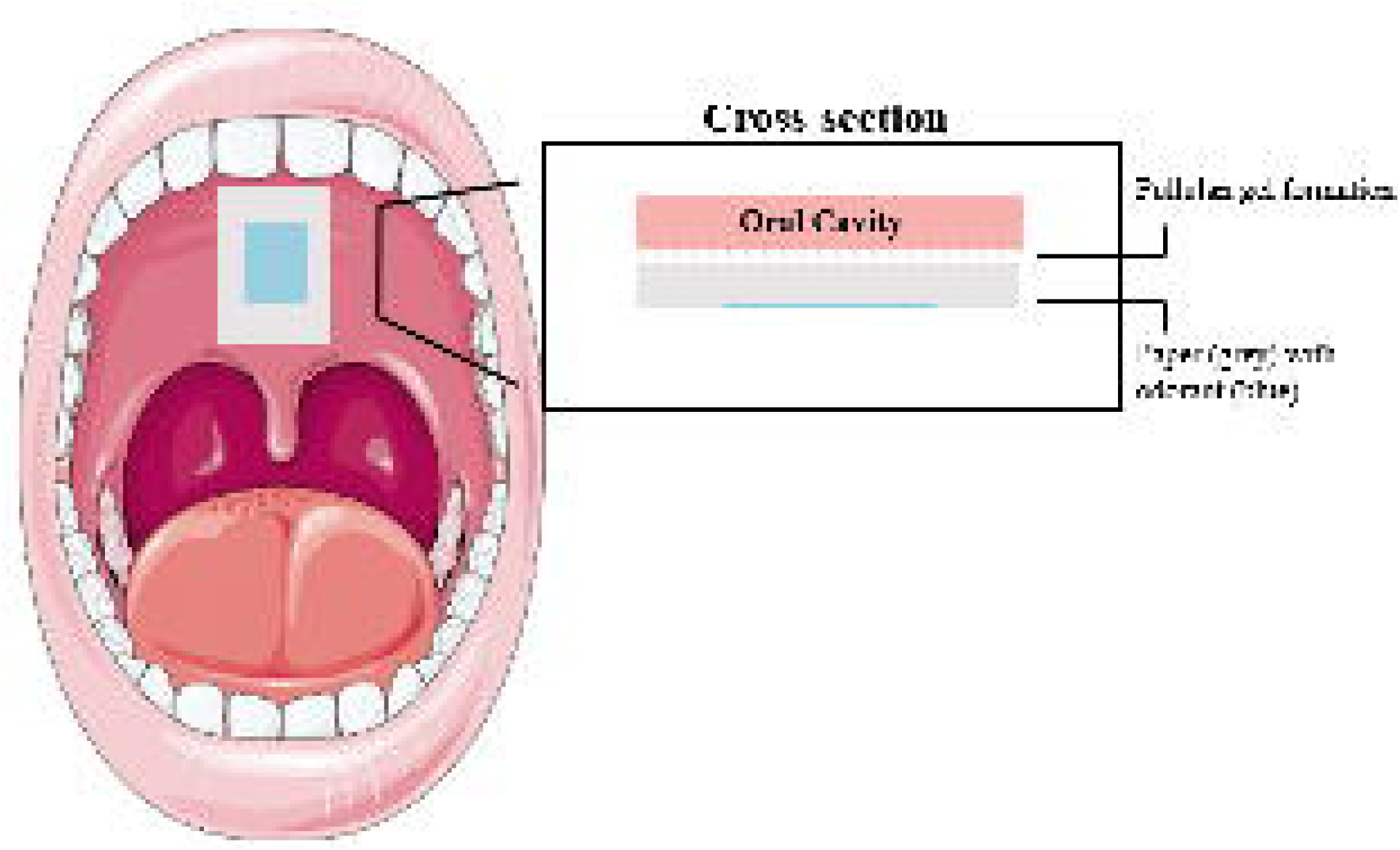

## Discussion

### Characterization of retronasal habituation

To date, only one study looking at habituation of retronasally perceived odors in humans has been performed (Pierce and Simons, 2018). In the present study, we demonstrated a temporal characterization of retronasal odor habituation in a naturalistic setting. We have confirmed the existence of retronasal odor habituation, outlined by two important principles of habituation laid out by Spencer and Thomson (1966), and later redefined by Rankin et al. (Thompson and Spencer, 1966; Rankin et al., 2010).

First, retronasal odors decline in strength after repetitive exposure thus following the underlying assumption of habituation set forth by Principle 1. This was measured with several parameters within the TI-curve including post peak area and time to extinction (T_ext_), and visually depicted in Figure 3. Similarly, these characteristics of habituation have been shown in numerous orthonasal odor studies (Cain, 1974; de Wijk, 1989; Schiet and Cain, 1990; Cain and Polak, 1992; Stuck et al., 2013; Sinding et al., 2017). However, most of these studies used single odorants and/or presented odors continuously through olfactometers. These studies are useful at characterizing habituation in lab settings, but are more difficult to translate to a real environment or situations like food/beverage intake. Many of these studies show faster habituation times for odorants than the data presented here, which may be a result of methodology differences or stimuli used. For instance, it has been shown that mixtures are more resistant to habituation than single odorants (Schiet and Cain, 1990; Berglund and Engen, 1993). Schiet and Cain point to an exceptional “durability” to habituation for odor mixtures – with some individuals not even completely habituating to some mixtures over extended exposure times (15 min).

It’s also important to note that in naturalistic settings, information enters the olfactory system in synchronized pulses following the respiratory system, unlike a continuous presentation by an olfactometer. Several animal models measuring neural activity have shown neurons within the piriform cortex are entrained to this cycle and that mitral/tufted cells, in the olfactory bulb (OB), tend to activate during inhalation and be inhibited during exhalation (Chaput and Panhuber, 1982; Chaput et al., 1992; Wilson, 1998). This activity, tied to respiration, likely reduces adaptation thus preventing complete habituation during normal breathing (Köster and de Wijk, 1991).

Secondly, the present study confirms that habituation is concentration-dependent for retronasally perceived odors, a characteristic defined by Principle 5. Using T-I curves, we showed that a lower lime concentration habituated faster and with a smaller post peak area, than a higher concentration. This result matches various experiments using other odorants presented orthonasally, either through respiratory action or an olfactometer, showing a larger reduction in odor intensity for lower concentrations compared to higher concentrations (Stone et al., 1972; Todrank et al., 1991; Cain and Polak, 1992; Stuck et al., 2013). An increase in physiological mechanisms (e.g. receptor recruitment) leading to an increased intensity perception has been an explanation for this effect, in which more receptors must adapt leading to a slower rate of habituation (Laing et al., 2003).

Lastly, we noted the non-food odor habituated slower than the food odor. Although lavender, an odor not typical of food, was perceived at the same intensity as lime it had a slower time to extinction and a larger post peak area. One explanation could be the context of odor perceived within the mouth, in which the perceiver is expecting a food-related odor and odors not fitting this description are perceived with caution. The considerable influence of cognitive processes on odor habituation has been demonstrated in several research studies (Dalton 1999; Dalton 1996; Dalton, Dilks, and Ruberte 1999; Kobayashi et al. 2007). In a series of experiments by Dalton and colleagues, odors given negative characterization prior to exposure, either through experimenter-provided instruction, verbal cues or behavior of a subject, showed much less habituation across exposure time. The route-dependency of the non-food related odor could be further evidence that non-sensory, top-down processes can delay retronasal olfactory habituation. Future research should focus on possible differences in retronasal habituation in food and non-food odors.

### Impact of habituation on flavor

In this study, we show in two separate experimental designs that habituation induced in naturalistic settings does not impact flavor perception. The fact that participants noticed no differences pre-post habituation to just lime oil dissolved in water points to at least partial simultaneous recovery. Several researchers have reported fast, almost immediate partial recovery, called simultaneous recovery, for orthonasal habituated odors (Cain, 1974; Cain and Polak, 1992; Gagnon et al., 1994; Philpott et al., 2008; Smith et al., 2010; Stuck et al., 2013). Retronasally presented odors are coupled with food consumption, so it is no surprise reports show that swallowing occurs significantly faster and more frequently under retronasal stimulation, compared to orthonasal stimulation (Welge-Lussen et al., 2009). This mechanical instinct, along with respiration as mentioned earlier, may contribute to dishabituation thus reducing the habituation effect and related recovery rates (Rankin et al., 2010).

Another explanation deals with compensation (de Wijk, 1989, Schiet and Cain, 1990; Berglund and Engen, 1993). Even though flavor is multisensory in nature, it is often perceived as a single sensory experience (Rozin, 1982). The absence of one part may be compensated by others (e.g. taste and odor) thus preserving the perceived intensity. A shift from an adapted to a non-adapted part has been shown within unimodal sensory systems such as vision (Foster, 2003) and even olfaction (Schiet and Cain, 1990; Berglund and Engen, 1993). For instance, when an individual adapts to a single odorant and then is presented a binary mixture, containing the adapted odorant, the overall odor intensity of the mixture stays relatively stable (Schiet and Cain, 1990; Berglund and Engen, 1993). Here, the perceived intensity of a weak component within a mixture may be unaltered because of prior exposure to another component.

### Limitations

This study was contrived in an exploratory manner, creating several limitations. We only used one food and one non-food odor, making any strong conclusions about possible differences in how we habituate to non-food versus food odor unreasonable. Additionally, a small delay was present between removal of odor stimuli and presentation of the flavor stimuli. We estimate this time to be approximately 3 sec on average, and could have influenced our attempts to characterize recovery. Lastly, while pullulan is widely considered tasteless and odorless, further studies would need to be done regarding odorant migration. It is possible, upon the formation of a pullulan hydrogel that adheres to the roof of the mouth, that odor molecules could migrate and come in contact with taste receptors. It is possible that if the odor stimuli used in this study contacted taste receptors, a bitter taste would be perceived. Additionally, taste stimulation could occur if a participant pressed on odor strip with their tongue. We don’t believe this to be a common occurrence, as a researcher was constantly observing the panelists perform the procedure and the maximum amount of odorant on any of the stimuli was only 3.3 μL. Taste stimulation would most likely modify our characterization of retronasal habituation.

## Conclusions

The studies presented here give clear characterization of the temporal changes in perception that take place with constant retronasal odor presentation. Furthermore, the aspect of this paper that may be of most importance to the food industry and nutrition professionals is that under *real-world* conditions, retronasal habituation does not appear to influence flavor perception. This finding was found in both simple and complex flavor stimuli, creating strong evidence for this conclusion. Transferring this finding to eating behavior, rapid and repetitive consumption of food/beverage is unlikely to influence flavor intensity of subsequent food intake. The findings of this study also give evidence that we habituate to different odors at different rates, more specifically we provide evidence that differentiates between food and non-food odors. Further research should explore this result with more rigor and the possible neural underpinnings of retronasal habituation to odors with different qualities.

